# Draft genome sequence of a *Candidatus* Brocadia bacterium enriched from tropical-climate activated sludge

**DOI:** 10.1101/123943

**Authors:** Xianghui Liu, Krithika Arumugam, Gayathri Natarajan, Thomas W. Seviour, Daniela I. Drautz-Moses, Stefan Wuertz, Yingyu Law, Rohan B. H. Williams

**Affiliations:** Singapore Centre for Environmental Life Sciences Engineering, Nanyang Technological University, 60 Nanyang Drive, SBS-01N-27, 637551, Singapore; School of Civil and Environmental Engineering, Nanyang Technological University, N1-01a-29, 50 Nanyang Ave, Singapore, 639798; Singapore Centre for Environmental Life Sciences Engineering, National University of Singapore, 21 Lower Kent Ridge Road 119077, Singapore

**Author notes:** X.H.L and K.A contributed equally to this work.

## Abstract

We present the draft genome of an anaerobic ammonium-oxidizing (anammox) bacterium, cluster III *Candidatus* Brocadia, which was enriched in an anammox reactor. A 3.2 Mb genome sequence comprising 168 contigs was assembled, in which 2,765 gene-coding regions, 47 tRNAs, and 5S, 16S and 23S ribosomal RNAs were annotated. No evidence for the presence of a nitric oxide-forming nitrite reductase was found.

---

Anaerobic ammonium oxidation (anammox) is a microbial bioprocess in which ammonium (NH_4+_) and nitrite (NO_2 −_) are converted to dinitrogen (N_2_) gas via the generation of hydrazine (N_2_H_4_) (*1*), and is considered a more energy efficient and effective N removal bioprocess than conventional wastewater treatment systems (*1*). All known anaerobic ammonia-oxidizing bacteria (AnAOB) belong to the phylum *Planctomycetes*, with five *Candidatus* genera defined (*1, 2*). We established an enrichment protocol to target anammox organisms in a sequencing batch reactor seeded with return activated sludge from a water reclamation plant (Public Utilities Board, Singapore). The reactor was fed synthetic wastewater containing ammonium and nitrite and operated at 35°C. Within six weeks, simultaneous ammonium and nitrite depletion was observed, along with the appearance of a reddish-brown biofilm on the walls of the reactor, highly characteristic of anammox biofilms (*1*). FISH analysis conducted on day 120 postinoculation using the Amx820 probe (*3*) indicated the presence of anammox bacteria. The wall biofilm was then sampled and shotgun metagenomics was performed on an Illumina MiSeq run using 300 bp paired end sequencing (Illumina). Following read filtering, we analysed these data using Ribotagger (*4*) and observed ∼70% relative abundance for genus *Candidatus* Brocadia (V4 ribotag annotated against SILVA release 119 (*5*)).

We constructed a metagenome assembly using Newbler v 2.9 (454 Life Sciences), and employed CONCOCT to cluster these contigs into 27 putative genome bins (*6*). The most abundant bin contained 168 contigs (length 1,527-114,900 bp; mean 19,210 bp) and had a total sequence length of 3.27Mb with an average %GC-content of 42.3%. Analysis with CheckM (*7*) showed this genome to have a high completeness (98.8%) and low contamination rate (1.7%), and was annotated to the marker set lineage UID2565, which is selective for *Candidatus* Brocadiaceae. We annotated predicted genes with Prokka (v1.10) (*8*), finding this draft genome encodes 2765 genes, 47 tRNAs, and 3 rRNA genes. Using the CheckM SSUfinder module (*7*), one 16S gene was identified from this bin, of length 1572bp, and which when analysed using SINA (*9*) was annotated to *Candidatus* Brocadia with 94% sequence identity. Collectively, these data suggest we have recovered a new member of genus *Candidatus* Brocadia.

This draft genome shows to encode gene clusters for core anammox metabolism including 9 multiheme *hdh/hao*-like proteins, the three subunits of *hzs* (hydrazine synthase), and a single, three sub-unit nitrate reductase/oxidoreductase (*nxr*) (*10*). The presence in anammox bacteria of nitrite reductases, either nitric oxide (NO)-forming and/or ammonia-forming, remains subject to considerable uncertainty (*10*). Using a sequence similarity network approach that incorporates all protein sequences from 1) the present metagenome assembly; 2) all extant draft genomes and 3) all known nitrate reductases, we identified a gene in the current draft genome that has a 88%-AAI and a 100% alignment overlap with ammonia-forming nitrite reductase in *Candidatus* Brocadia sinica JPN1 (E=1x10^-257^), and at a lower level of support, with an ammonia-forming nitrite reductase in genus *Rhodopirellula* (53% AAI, 94% alignment overlap, E=1×10^-146^). We did not find any clear evidence for the presence of a NO-forming *nir* gene, consistent with previous investigations of the genus Brocadia (10-13).

## Accession number(s)

The draft *Candidatus* Brocadia genome from this study was deposited in the NCBI Genome under the submission number SUB2595315 and the raw sequence data from the reactor community is available via NCBI BioProject Accession PRJNA383778.

## ACKNOWLEDGMENTS

This research was supported in part by the Singapore National Research Foundation and the Ministry of Education under the Research Centre of Excellence Programme and in part by the Singapore National Research Foundation grants 1102-IRIS-10-03 and 1301-IRIS-59. We thank Nguyen Thi Quynh Ngoc for performing DNA extraction, Larry Liew for assistance with sampling, Sara Swathi and Eganathan Kaliyamoorthy for laboratory measurements and Dr Yeshi Cao, Azhar Kasmani, Roslan Dawi, Hui Yui, Bee Hong Kwok, Dr Winson Lay and Dr Yangshuo Gu of the Public Utilities Board for their assistance and collaboration.

## References

1. Kartal, B., de Almeida, N.M., Maalcke, W.J., Op den Camp, H.J., Jetten, M.S., Keltjens, J.T. 2013. How to make a living from anaerobic ammonium oxidation. FEMS Microbiol Rev. 37: 428–461.

2. Oshiki, M., Satoh, H., Okabe, S. 2015. Ecology and Physiology of Anaerobic Ammonium Oxidizing Bacteria. Environ Microbiol. 18: 2784–2796.

3. Daims H, Hilde L, Nielsen PH. 2009. FISH Handbook for Biological Wastewater Treatment: Identification and Quantification of Microorganisms in Activated Sludge and Biofilms by FISH. International Water Association, London.

4. Xie, C., Goi, C.L.W., Huson, D.H., Little, P.F.R., Williams, R.B.H. 2016. Ribtoagger: fast and unbiased 16S/18S profiling using whole community shotgun metagenomic or metatranscriptome surveys, BMC Bioinformatics 17(Suppl 19): 508.

5. Pruesse, E., Quast, C.,Knittel, K., Fuchs, B. M., Ludwig, L., Peplies, J., Glöckner, F. O. 2007. SILVA: a comprehensive online resource for quality checked and aligned ribosomal RNA sequence data compatible with ARB. Nucleic Acids Research 35: 7188–7196.

6. Alneberg, J., Bjarnason, B. S., de Bruijn, I., Schirmer, M., Quick, J., Ijaz, U. Z., Lahti, L., Loman, N.J., Andersson, A.F., Quince, C. 2014. Binning metagenomic contigs by coverage and composition. Nat Methods. 11: 1144–1146.

7. Parks, D.H., Imelfort, M., Skennerton, C.T., Hugenholtz, P., Tyson, G.W. 2015. CheckM: assessing the quality of microbial genomes recovered from isolates, single cells, and metagenomes. Genome Res. 25:1043–1055.

8. Seemann, T. 2014. Prokka: rapid prokaryotic genome annotation. Bioinformatic. 30: 2068–2069.

9. Pruesse, E., Peplies, J., Glöckner, F.O. 2012. SINA: accurate high-throughput multiple sequence alignment of ribosomal RNA genes. Bioinformatic. 28: 1823–1829.

10. Kartal, B. and Keltjens, J. T. 2016. Anammox biochemistry: a tale of heme c proteins, Trends in Biochemical Sciences 41(12): 998–1011.

11. Oshiki, M., Shinyako-Hata, K., Satoh, H., Okabe, S. 2015. Draft genome sequence of an anaerobic ammonium-oxidizing bacterium, “*Candidatus* Brocadia sinica.” Genome Announc 3(2): e00267–15.

12. Ali, M., Haroon, M. F., Narita, Y., Zhang, L., Shaw, D. R., Okabe, S., Saikaly, P. E. 2016. Draft genome sequence of the anaerobic ammonium-qxidizing bacterium “*Candidatus* Brocadia sp. 40”. Genome Announc. 4 (6): e01377–16.

13. Speth, D. R., in’t Zandt, M. H., Guerrero-Cruz, S., Dutilh, B. E., Jetten, M. S. M. 2015. Genome-based microbial ecology of anammox granules in a full-scale wastewater treatment system, Nature Communications 7: 11172

